# Transient Receptor Potential Ankyrin 1 Mediates Afferent Signals in the Inflammatory Reflex

**DOI:** 10.1101/822734

**Authors:** Harold A. Silverman, Manojkumar Gunasekaran, Eric H. Chang, Qing Chang, Meghan E. Addorisio, Andrew Stiegler, Adam M. Kressel, Jian Hua Li, Tea Tsaava, Tomas S. Huerta, Valentin A. Pavlov, Ulf Andersson, Sangeeta S. Chavan, Kevin J. Tracey

**Affiliations:** Center for Biomedical Science and Bioelectronic Medicine, Feinstein Institutes for Medical Research, Northwell Health, 350 Community Drive, Manhasset, NY 11030, USA; Donald and Barbara Zucker School of Medicine at Hofstra/Northwell, 500 Hofstra University, Hempstead, New York 11030, United States of America; Department of Surgery, North Shore University Hospital, Northwell Health, 300 Community Drive, Manhasset, NY 11030, USA; The Elmezzi Graduate School of Molecular Medicine, 350 Community Drive, Manhasset, NY 11030, USA; Department of Women’s and Children’s Health, Karolinska Institute, Karolinska University Hospital, 17176 Stockholm, Sweden

## Abstract

Survival of an organism requires mechanisms to sense damaging factors in the environment. In mammals, bacterial toxins and inflammatory mediators stimulate nociceptive sensory neurons to activate protective reflexes. Whereas the vagus nerve reflex circuit that protects against damaging inflammation, termed the “inflammatory reflex,” was described more than twenty years ago^1,2^, how the vagus nerve detects inflammation to initiate the inflammatory reflex has remained unknown. Here we show that transient receptor potential ankyrin 1 (TRPA1) in sensory vagus neurons is required to sense interleukin-1β (IL-1β), a central cytokine mediator of inflammation and injury. Selective activation of vagus nerve TRPA1 using optopharmacology stimulated the inflammatory reflex to inhibit innate inflammatory responses to bacterial lipopolysaccharide and IL-1β. Proximity ligation assay and immunohistochemistry revealed that IL-1 receptors are co-expressed with TRPA1 in vagus sensory neurons. Whole-cell patch-clamp recordings reveal that TRPA1 is required to mediate IL-1β-dependent depolarization of vagus sensory neurons. Further, TRPA1-deficient mice lack inflammatory reflex attenuation of inflammation, fail to restrain cytokine release, and have significantly enhanced lethality to bacterial sepsis. Therefore, vagus neurons expressing TRPA1 are necessary and sufficient to activate the sensory arc of the inflammatory reflex to protect against harmful inflammation.

---

Reflex neural circuits form the basis for physiological responses when the body is exposed to changing environmental and physiological conditions. The vagus nerve, originating in the brainstem medulla oblongata, is a paired structure that travels through the neck, thorax and abdomen to innervate visceral organs. In humans it is comprised of approximately 100,000 neurons; 80% are sensory, transmitting information to the brainstem in response to changes in the body’s internal milieu to stimulate reflex motor signals that regulate heart rate, digestion, and other organ functions. The inflammatory reflex, a vagus circuit that inhibits inflammation to protect tissues from damage, stimulates anti-inflammatory T cells expressing choline acetyltransferase (T ChAT) in the spleen to produce acetylcholine which interacts with α7 nicotinic acetylcholine receptor (α7nAChR)-expressing macrophages to inhibit release of TNF, IL-1, HMGB1 and other inflammatory cytokines^1–3^. Electronic stimulation of the inflammatory reflex confers significant protection in preclinical models of arthritis, inflammatory bowel disease, sepsis, and other syndromes, and significantly improves outcomes in clinical trials of rheumatoid arthritis and Crohn’s disease^2,4,5^. Despite a detailed mechanistic understanding of signaling in the motor arc of the inflammatory reflex, the afferent arc mechanisms were previously unknown. TRPA1, a polymodal nociceptor cation channel expressed by unmyelinated peptidergic nociceptive C fiber neurons, is stimulated by reactive (hydrogen peroxide, formalin, acrolein) and non-reactive (nicotine) compounds^6,7^. TRPA1 is also activated by proinflammatory molecules, bacterial toxins, reactive oxygen and nitrogen species, bradykinin, prostaglandins, bacterial products, and environmental and chemical irritants^8–10^. Because TRPA1 nociceptors are expressed in vagus neurons, we reasoned that TRPA1 might be a candidate sensor responding to inflammation in an afferent arc of the inflammatory reflex.

We utilized optopharmacology to directly stimulate vagus nerve TRPA1 nociceptors in mice receiving bacterial endotoxin (lipopolysaccharide, LPS), a potent stimulus to cytokine production and inflammation. We observed that stimulation of TRPA1-expressing vagus fibers using optovin, a photosensitive TRPA1 ligand^11^, significantly inhibited serum TNF levels in wild-type mice during endotoxemia as compared to sham stimulation (Fig. 1a). Optovin-mediated photostimulation of the vagus nerve failed to reduce TNF levels in TRPA1-deficient mice, giving direct evidence that vagus nerve TRPA1 activation inhibits TNF (Fig. 1b). To determine the directionality of optovin-mediated vagus TRPA1 signal transmission, we carried out selective vagotomy. Cutting the vagus nerve rostral to the optically-stimulated area, thus eliminating afferent vagus nerve signals to the brainstem while maintaining efferent vagus signals to the spleen, abrogated TRPA1-dependent inhibition of TNF (Fig. 1a and 1c). In contrast, vagotomy caudal to the stimulation site failed to abolish suppression of TNF after photochemical stimulation of the vagus nerve (Fig. 1c). To address whether activation of vagus TRPA1 nociceptors stimulates the inflammatory reflex, α7nAChR-deficient mice were subjected to optovin photostimulation and endotoxemia. TRPA1-dependent activation of the afferent vagus nerve signals failed to attenuate serum TNF in endotoxemic α7nAChR-deficient mice (Fig. 1d) indicating that TRPA1-mediated inhibition of TNF requires α7nAChR. Together with prior results that the inflammatory reflex requires α7nAChR to inhibit TNF production^12^, these observations indicate that vagus TRPA1 nociceptors are necessary and sufficient to stimulate the afferent arc of the inflammatory reflex.

**Fig. 1.**
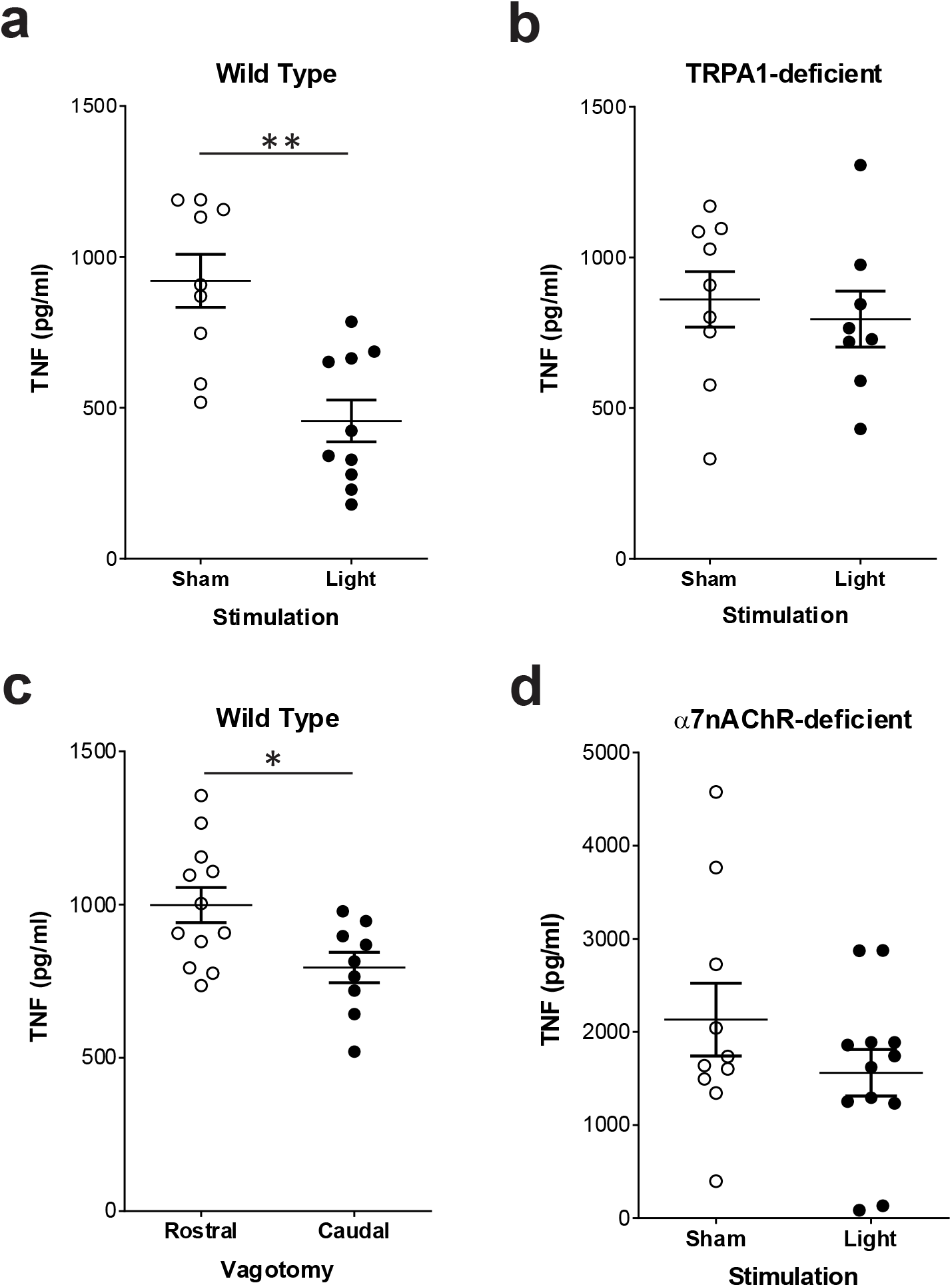
Optovin photostimulation of the vagus nerve requires TRPA1 to attenuate TNF in endotoxemia. Optovin was directly applied to the cervical vagus nerve and exposed to sham or 405 nm light stimulation prior to endotoxin injection. Blood was obtained 90 min after endotoxin administration, and serum TNF was measured by ELISA. **a**, wild-type mice receiving sham (n=9) or light (n=10) stimulation. **b**, TRPA1-deficient mice receiving sham (n=9) or light (n=8) stimulation. **c**, wild-type mice were subjected to rostral (n=12) or caudal (n=9) vagotomy followed by optovin stimulation. **d**, α7nAChR-deficient mice receiving sham (n=10) or light (n=10) stimulation. Mean ± SEM. *p<0.05; **p<0.01 (Mann-Whitney test).

Proximity ligation assay and immunohistochemical analysis reveal that TRPA1 is colocalized with IL1-receptor1(IL1R1) on sensory vagus neurons isolated from nodose ganglia, where cell bodies of vagus sensory afferents reside (Fig. 2a-b). TRPA1 is a non-selective cation channel that mediates calcium uptake and action potentials, which can be studied using intracellular calcium imaging and electrophysiology^7^. To assess IL-1β responsiveness of TRPA1-expressing sensory vagus neurons, we studied intact nodose ganglion isolated from Vglut2-GCaMP3 mice, in which the genetically-encoded Ca^2+^ indicator GCaMP3 is specifically expressed in >99% of nodose sensory neurons under the control of *Vglut2* promoter^13,14^. IL-1β induces significant calcium influx in a subset of sensory vagus neurons that also respond to polygodial, a TRPA1 agonist^15^ (Supplemental Data Fig. 1, Table 1). Most TRPA1-expressing nociceptor neurons co-express the transient receptor potential vanilloid 1 (TRPV1) receptor^16,17^. Accordingly, we observe that more than 90% of the polygodial-responsive neurons in the nodose ganglia also respond to capsaicin, a potent TRPV1 agonist^18^ (Supplemental Data Fig. 1, Table 1). By tracking GCaMP3 activation, we identified neurons residing in nodose ganglia that selectively respond to IL-1β, capsaicin (TRPV1-positive) and polygodial (TRPA1-positive) (Supplemental Data Fig. 1, Table 1). Using Fluo-4 based *in vitro* calcium imaging we confirmed that IL-1β also activated isolated nodose sensory neurons (Fig. 2c). Moreover, we observed that most IL-1β-responsive neurons from wild-type nodose ganglia also responded to polygodial (Fig. 2c, Table 1). In contrast, nodose ganglion neurons isolated from TRPA1-deficient mice fail to respond to IL-1β (0/36) and polygodial (0/36) but maintain sensitivity to the TRPV1 agonist capsaicin (36/36) (Fig. 2c, Table 1). We also evaluated the effect of the selective TRPA1 antagonist AM0902^19^, and observed that it inhibits IL-1β-induced calcium responses in TRPA1-expressing nodose sensory neurons (Fig. 2d and Supplemental Data Fig. 2).

**Fig. 2.**
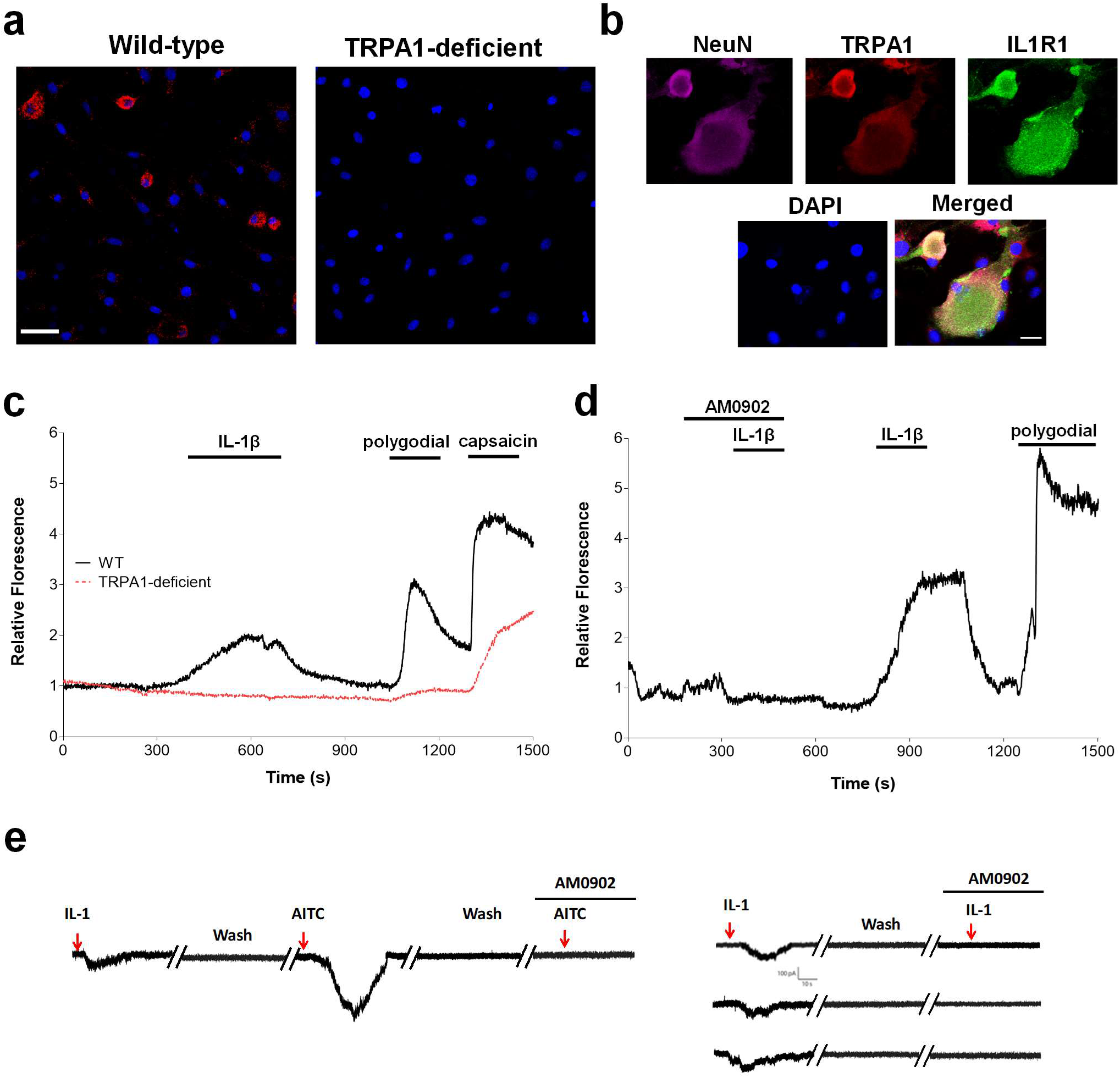
TRPA1 and IL1R1 are within close proximity, and TRPA1 is required for IL-1β-mediated activation of nodose sensory neurons. **a**, Representative *in situ* image of proximity ligation assay showing close proximity of TRPA1 and IL1R1 on a subset of nodose ganglion neurons (Scale bar = 50 μm). Nodose sensory neurons were isolated from wild-type and TRPA1-deficient mice and subjected to *in situ* proximity ligation assay (Methods), using primary antibody pairs of anti-TRPA1 and anti-IL1R1 antibodies. Dual binding by a pair of corresponding proximity probes, secondary antibodies with attached oligonucleotides, generates a spot (red colored blob) if the two antibodies are in close proximity. Thus, each individual blob represents TRPA1 in close proximity with IL1R1. The cells were counterstained with DAPI (blue) to visualize the nucleus. **b**, Representative images of immunohistochemical staining of cultured nodose ganglion neurons isolated from wild-type mice to detect neurons (NeuN, magenta), TRPA1 (red), IL1R1 (green), and nuclei (DAPI, blue). 400X magnification; scale bar = 10μm. **c**, Example of the effect IL-1β (20 μg/ml) on [Ca^2+^]_i_ levels in nodose ganglion neurons isolated from wild-type mice (black solid line) and TRPA1-deficient mice (red dashed line). Polygodial (10 μM) and capsaicin (10 μM) are applied to identify TRPA1- and TRPV1-expressing neurons, respectively. **d**, Representative GCaMP3 fluorescence signal in cultured nodose ganglia neurons from Vglut-GCamp3 mice showing responses to IL-1β (20 μg/ml) in the presence and absence of TRPA1 antagonist AM0902 (10 μM). Polygodial (10 μM) is applied to identify TRPA1-expressing neurons, **e**, Responses to IL-1β and TRPA1 agonists AITC monitored with whole-cell patch-clamp recordings in isolated nodose ganglion neurons. When held at −70 mV, IL-1β (20 μg/ml) and AITC (100 μM) induce inward currents which are blocked by bath application of TRPA1 antagonist AM0902 (50 μM) (top tracing). Three additional IL-1β-responsive neurons are shown below, with AM0902 blocking inward currents in all experiments.

**Table 1.** **Percentage breakdown of responsive nodose ganglia sensory neurons** that respond to polygodial, IL-1β and both IL-1β and polygodial in intact nodose ganglia and cultured nodose ganglion neurons isolated from Vglut-GCamp3, wild-type and TRPA1-deficient mice.

| Prep | Strain | Capsaicin | Polygodial | IL-1β | Polygodial + IL-1β |
| --- | --- | --- | --- | --- | --- |
| Intact | VGlut2-GCamp3 | 100%<br>(225/225) | 23.9%<br>(61/225) | 4.3%<br>(11/225) | 3.5%<br>(9/225) |
| Cultured | Wildtype | 100%<br>(138/138) | 69.1%<br>(94/138) | 2.2%<br>(3/138) | 2.2%<br>(3/138) |
| Cultured | TRPA1-deficient | 100%<br>(36/36) | 0.0%<br>(0/36) | 0.0%<br>(0/36) | 0.0%<br>(0/36) |

Previous studies have shown that IL-1β increases the excitability of dorsal root ganglia nociceptors^20^. Therefore, we measured IL-1β-induced currents in whole-cell patch-clamp recordings in a subset of nodose ganglion sensory neurons that were first screened for IL-1β responses using calcium transient analysis. When the neuron membrane potential is held at −70 mV, application of IL-1β (20μg/ml) produces slow inward currents in 4 out of 332 (1.2%) nodose sensory neurons in culture (Fig. 2e). The mean amplitude of IL-1β induced currents is 108 ± 23 pA (n = 4). Neurons responding to IL-1β also respond to allyl isothiocyanate (AITC, 100 μM; Fig. 2e), a selective TRPA1 activator^9^. The peak current (410 ± 39 pA, n = 4) induced by AITC is approximately 3.8-fold greater than that elicited by IL-1β (Fig. 2e). Application of selective TRPA1 antagonist, AM0902 (50 μM), together with IL-1β significantly prevents the activating effects of IL-1β (n = 3; Fig. 2e). In current-clamp mode, the threshold of action potentials (−25 ± 2 mV) induced by current injections (100 pA, 200 ms) is reduced by IL-1β application to −31 ± 3 mV (n = 4), but is reversed fully within 30 mins after washout (data not shown). Thus, IL-1β-dependent increases in membrane excitability require TRPA1-expressing sensory neurons.

In agreement with our recent results^21,22^, administration of IL-1β to wild-type mice induces afferent vagus neuronal signal propagation (Fig. 3a, 3c, and Supplemental Data Fig. 3). No change in vagus nerve activity is observed in TRPA1-deficient mice following IL-1β administration (Fig. 3b-c, and Supplemental Data Fig. 3), indicating that TRPA1 is required for IL-1β–mediated sensory vagus signal transmission. To determine the role of TRPA1 in mediating the inflammatory responses to IL-1β, we administered IL-1β intraperitoneally to wild-type and TRPA1-deficient mice, and measured serum cytokine levels and body temperature. We observed that serum and spleen IL-6 and CXCL-1 levels are significantly increased in mice devoid of TRPA1 as compared to wild-type mice (Fig. 3d-e and Supplemental Data Fig. 4-5).

**Fig. 3.**
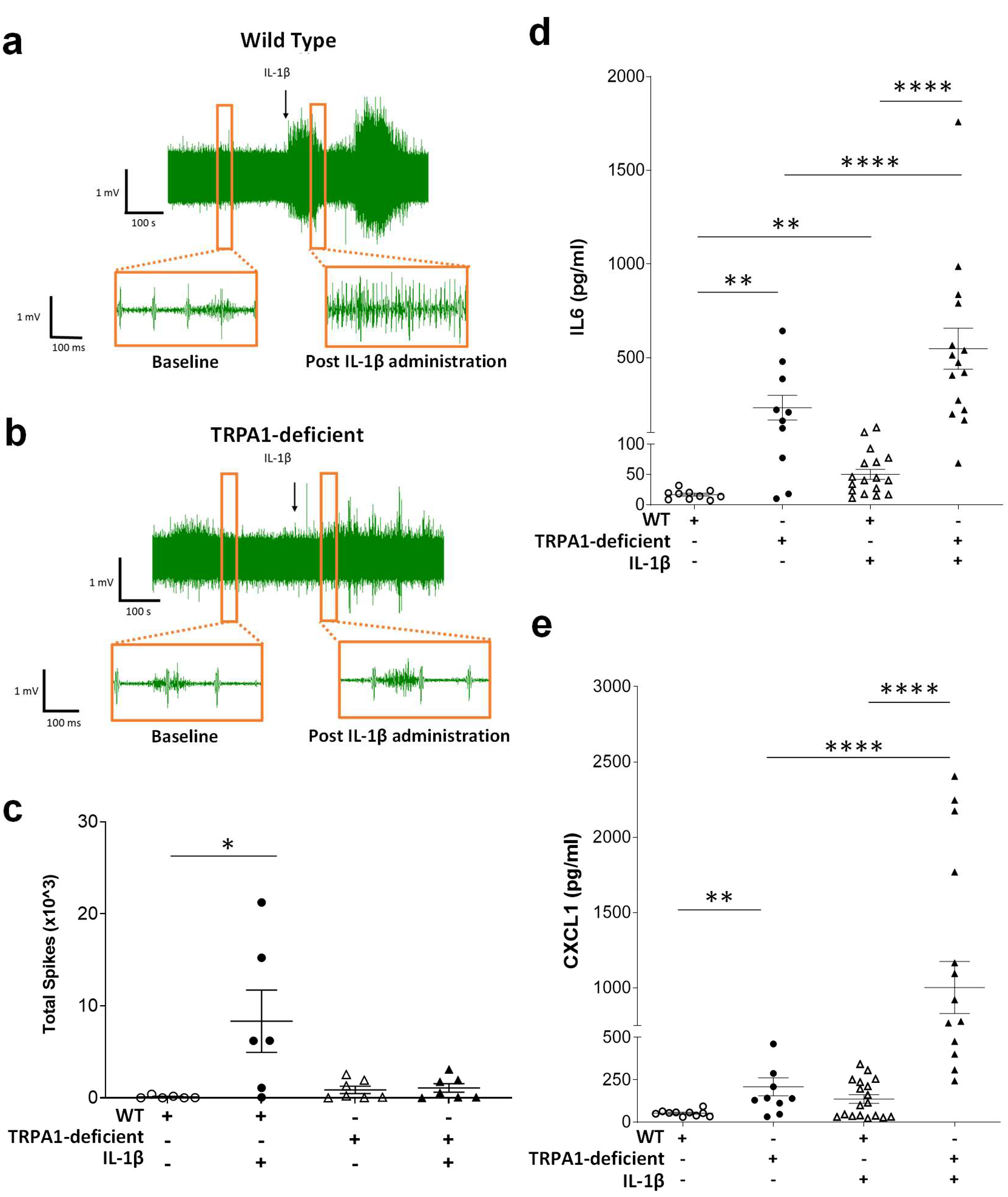
IL-1β-administration increases vagus nerve activity and circulating cytokine levels. Representative recordings of the vagus nerve signals from **a**, wild-type mice and **b**, TRPA1-deficient mice starting 2.5 min before (baseline), and 2.5 min after IL-1β administration (350 ng/kg) (upper traces). An enhanced view of the baseline (left inset), and post injection responses (right inset) is shown (lower traces). **c**, Total spike count over the entire 5-minute pre- and 5-minute post- IL-1β administration recordings in wild-type (n = 6) (open and close circles) and TRPA1-deficient mice (n = 7) (open and close triangles). Mean ± SEM. *p<0.05 (Paired t-test). Serum levels of **d**, IL-6 and **e**, CXCL1 in wild-type and TRPA1-deficient mice injected intraperitoneally with either saline (n = 9-10 per group) or IL-1β (3.2 μg/kg, n = 17-18 per group). Mean ± SEM. **p<0.01; ****p<0.0001 (Mann-Whitney test).

Administration of IL-1β significantly increases cytokine levels in wild-type mice, and these increases are even significantly higher in TRPA1-deficient mice as compared to wild-type mice (Fig. 3d-e and Supplemental Data Fig. 4-5). As expected from prior studies of sickness behaviour^23^, administration of IL-1β causes hypothermia in wild-type mice housed at an ambient temperature of 22°C (Fig. 4a). TRPA1-deficient mice, however, fail to develop hypothermia after administration of IL-1β (Fig. 4a) indicating that TRPA1 is required to mediate the cytokine and thermoregulatory responses to IL-1β. Because the inflammatory reflex is an evolutionarily conserved anti-inflammatory mechanism to protect against harmful inflammation, we next reasoned that a defective inflammatory reflex in TRPA1-deficient mice would render animals increasingly sensitive to sepsis. Wild-type and TRPA-deficient mice were subjected to non-lethal cecal-ligation and puncture (CLP) induced sepsis, and disease severity and mortality monitored. Sepsis severity is significantly increased within three days in TRPA1-deficient mice as compared to wild-type mice (p<0.0001, Fig. 4b). Wild-type mice significantly improved within 6 days and survived. TRPA1-deficient mice have persistently severe disease scores (Supplemental Data Fig. 6), and while none of the wild-type mice died following CLP, mortality is significantly increased in TRPA1-deficient mice (25% mortality, p<0.05, Fig. 4c).

**Fig. 4.**
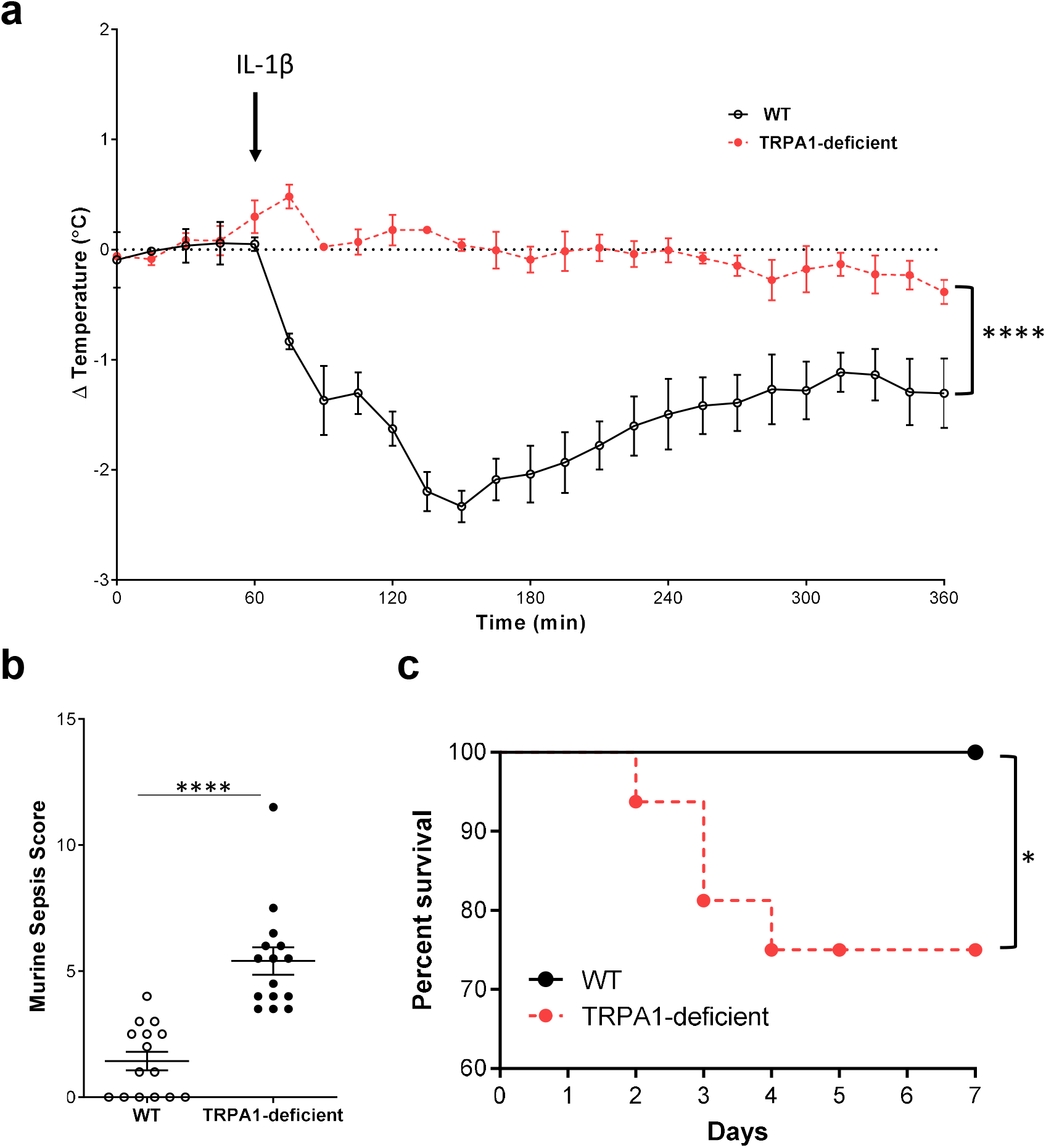
TRPA1 is required for the thermoregulatory and inflammatory responses to IL-1β. **a**, Core body temperature in wild-type (n = 4) and TRPA1-deficient (n = 4) mice was recorded for 60 min before and 300 min after intraperitoneal IL-1β (5.0 μg/kg) administration. Data were recorded at 15-min intervals and are presented as mean ± SEM. ****p<0.0001 IL-1β -injected wild-type mice vs. TRPA1-deficient mice (two-way ANOVA). **b**, Murine Sepsis Score on post-CLP sepsis day 3 in wild-type and TRPA1-deficient mice. Mean ± SEM. ****p<0.0001 (Mann-Whitney test) comparison between wild-type and TRPA1-deficient mice (n = 15 per group). **c**, Sepsis mortality increases in TRPA1-deficient mice. Non-lethal CLP causes no mortality in wild-type mice but 25% mortality in TRPA1-deficient mice. *p<0.05 (Log-rank test, n = 10 per group).

The major findings of the current study establish that vagus TRPA1 nociceptors are necessary and sufficient for IL-1β-mediated activation of the afferent arc of the inflammatory reflex. TRPA1 is co-expressed with IL-1β receptors on vagus sensory neurons, mediates IL-1β-induced vagus sensory neuron activation, and is required for the thermoregulatory and cytokine responses to IL-1β. TRPA1-deficiency impairs the inflammatory reflex control of inflammation and enhances the lethality of sepsis. It appears that vagus TRPA1 nociceptors have a proximate role in the thermoregulatory, behavioral, and inflammatory responses to infection. Although originally studied as a therapeutic target for pain, itch and respiratory disease, recent evidence suggests a broader role of TRPA1 as a sensor for inflammatory mediators^7–9^. Ablation of TRPV1-lineage neurons, which include the subpopulation of TRPA1-expressing nociceptors^17^, also enhances neutrophil infiltration, proinflammatory cytokine production, and adhesion molecule expression during endotoxemia^24^. Early work by Linda Watkins implicated vagus sensory neurons in mediating the sickness behavior responses to IL-1β^25^. Together with the established role of the inflammatory reflex in inhibiting cytokine release and alleviating inflammation^1–5^, the identification here of TRPA1 as a proximal mechanism provides a new insight into the role of sensing inflammation to stimulate protective inflammatory reflexes. Although these findings do not exclude the possibility that other nociceptors stimulated by other molecules may also activate the inflammatory reflex, impairment of TRPA1 afferent signaling pathways in the presence of IL-1β can lead to increased damage during inflammation or infection. These findings give new mechanistic insights into the fundamental intersection of neuroscience and immunity.

## METHODS

### Animals

All animal experiments were performed in accordance with the National Institutes of Health Guidelines under protocols approved by the Institutional Animal Care and Use Committee (IACUC) and the Institutional Biosafety Committee (IBC) of the Feinstein Institute for Medical Research, Northwell Health, Manhasset, New York, USA. Male B6.129F, TRPA1-deficient (B6;129P-*Trpa1^tm1Kykw^*/J), α7nAChR-deficient (B6.129S7-Chrna7 tm1Bay /J), TRPV1-cre (B6.129-*Trpv1^tm1(cre)Bbm^*/J), Vglut2-ires-cre (*Slc17a6^tm2(cre)Lowl^*/J), and GCaMP3 (B6.Cg-*Gt (ROSA)26 Sor^tm38(CAG-GCaMP3)Hze^*/J) mice were purchased from Jackson Labs. Animals were allowed to acclimate for at least five days prior to initiating the experiment. GCaMP3 was selectively expressed in glutamatergic neuron cell bodies using a Cre driver line (*Vglut2-ires-Cre*) that targets all vagus sensory neurons^13,14^ and a Cre-dependent reporter allele (*lox-GCaMP3*). Mice were housed under reverse day/light cycle and had access to food and water *ad libitum*. Food was withheld for the 3-4 hours prior to nerve recording; animals continued to have access to water.

### Surgical isolation of cervical vagus nerve

Mice were induced with general anesthesia using isoflurane at 2.5% in 100% oxygen at a flow rate of 1 L/min. Mice were then placed in the supine position and maintained at 2% isoflurane during surgery. The core body temperature was monitored with a rectal probe and was maintained around 37°C with a heating pad and heat lamp. The vagus nerve was then exposed and placed on the recording electrode as described previously^22,26^.

### Optopharmacological stimulation of the vagus nerve

Mice were induced with general anesthesia using isoflurane at 2.5% in 100% oxygen at a flow rate of 1 L/min and maintained in supine position at 2.0% isoflurane. The left vagus nerve was surgically exposed as described previously^22,26^.Optovin (2 μl of 15 mM) was directly applied on the nerve 2 minutes prior to light stimulation. Animals were subjected to 405 nm light stimulation (1000 mA, 10 hz, with a 10% duty cycle for 5 minutes with an approximate power of 80-85 μW) using Thor Labs LED driver DC4100, with a 405 nm LED model M405L3 (Newton, New Jersey). Sham stimulation group underwent similar surgical procedure and optovin application but no light stimulation. After 2 hr, animals were challenged with LPS (*Escherichia coli* 0111:B4, Sigma; 8 mg/kg, intraperitoneal administration). Animals were euthanized after 1.5 hr and blood was collected. Serum TNF levels were quantitated using commercial ELISA (R&D Systems).

### Vagotomy procedure

Prior to the vagotomy a 6.0 suture was tied around the vagus nerve, and a surgical cut of the vagus nerve was completed either rostral or caudal to the optovin application site, using the brain as the point of reference. Animals were then subjected to optovin application, light stimulation and endotoxin administration as described earlier. Mice were euthanized 1.5 hr after LPS administration, and blood was collected for TNF quantitation.

### Isolated sensory neurons

Nodose ganglia from B6.129F, TRPA1-deficient or *Vglut2*-Cre^+^.*R26*^LSL-GCamp3/+^ were harvested into ice-cold neurobasal medium (Gibco, Thermo Fisher Scientific, Waltham, MA, USA), dissociated with 1 μg/mL collagenase/dispase (Roche Life Science, Germany) in neurobasal medium for 90 minutes at 37°C on a rotator-shaker. After digestion, the nodose ganglia were washed with HBSS (Gibco), triturated with fire-polished glass Pasteur pipettes of decreasing size (Fisher Scientific, Waltham, MA, USA) and filtered through a 70 μM strainer. The cells were plated on poly-D-lysine and laminin coated glass cover slips in 24 well tissue culture plate in complete neurobasal medium [Neurobasal™ medium supplemented with penicillin-streptomycin (Gibco), GlutaMax™ (Gibco), B-27® serum-free supplement (Gibco), 50 ng/ml NGF (Sigma- Aldrich), and used for proximity ligation assay, immunohistochemical analysis, intracellular calcium measurements and whole-cell patch-clamp recordings within 24-48 h post plating.

### Proximity ligation assay (PLA)

The interactions between TRPA1 and IL-R1 in nodose ganglia cultures were detected by PLA with Duolink *in situ* kit (Sigma- Aldrich). Primary nodose ganglion cells were harvested and cultured on glass coverslips as described above. The cells were fixed with 4% paraformaldehyde/PBS for 15 min, washed with PBS, permeabilized with 0.2% Triton X-100 for 15 min, blocked with Duolink blocking solution for 60 min at 37°C, and then incubated in primary antibodies diluted in Duolink antibody diluent overnight at 4°C. Primary antibodies were rabbit anti-TRPA1 (polyclonal, 1:500; Millipore, Temecula, CA) and goat anti-IL R1 (polyclonal, 1:40; R&D Systems). After incubation with primary antibodies, the cells were washed, incubated with Duolink anti-goat PLUS and anti-rabbit MINUS PLA probe solutions at 37°C in. After 60 min, cells were washed and incubated with the ligation solution for 30 min at 37°C. After ligation, the cells were washed and incubated with amplification solution for 100 min at 37°C. After washing, cells were mounted using Duolink mounting medium with DAPI. Labeled cells were visualized and imaged using a confocal microscope (Zeiss). The resulting positive signals were recognized as discrete fluorescent spots. Each spot represents one interaction event. All specimens were imaged under identical conditions and analyzed using identical parameters.

### Immunohistochemistry

Primary nodose ganglion cells were harvested and cultured on glass coverslips as described above. The cells were fixed with 4% paraformaldehyde/PBS for 15 min, washed with PBS, permeabilized with 0.2% Triton X-100, blocked with 10% donkey serum, and then incubated with a cocktail of primary antibodies diluted in PBS containing 2% donkey serum and 0.05% Triton X-100 overnight at 4 °C. The primary antibodies were: rabbit anti-TRPA1 (polyclonal, 1:500; Millipore, Temecula, CA); goat anti-IL1R1 (polyclonal, 1:40; R&D Systems, Minneapolis, MN); and mouse anti-NeuN (monoclonal, 1:50; Millipore). After incubation with primary antibodies, the cells were washed four times with PBS, then incubated for 2 hr at room temperature in a mixture of species-specific secondary antibodies (all raised in donkey) conjugated to Alexa Fluor 488, Alexa Fluor 594 and Alexa Fluor 647 (Invitrogen Corporation, Carlsbad, CA). Coverslips were washed again and mounted using anti-fade mounting medium with DAPI (Vectashield; Vector Laboratories, Burlingame, CA).

### Calcium imaging

Sensory nodose neurons isolated from B6.129F and TRPA1-deficient mice were loaded with Flou-4 NW with pluronic acid F-127 (Molecular Probes) for 60 minutes at 37°C for 45 min in neurobasal-A medium, washed, and imaged at room temperature. For imaging the nodose neurons from Vglut2-GCaMP3 mice, cells were isolated, cultured on coverslips as described above and imaged. Confocal images were acquired continuously at a frame rate of 4 Hz with an imaging resolution of 512 × 512 pixels while chemical agonists (20 μg/ml IL-1β, 10 μM polygodial and 10 μM capsaicin) applied *via* the valve-controlled perfusion system on a Zeiss LSM-880 confocal laser microscope (Warner Instruments). AM0902 was perfused as indicated in respective experiments. Cells were washed with HBSS before application of each new agonist. Individual neurons were identified and analyzed for fluorescence intensity changes offline.

### Nodose ganglion imaging

For *ex vivo* imaging, intact nodose ganglia along with a part of the cervical vagus nerve were excised from Vglut2-GCaMP3 mice, placed into HBSS buffer, and mounted into a custom-fabricated glass holding chamber. This chamber is designed to sit within a standard liquid perfusion system (Warner Instruments) mounted onto the imaging stage of a Zeiss LSM-880 confocal laser microscope. Confocal images were acquired continuously at a frame rate of 4 Hz with an imaging resolution of 512 × 512 pixels while chemical agonists were applied via the valve-controlled perfusion system (Warner Instruments). Neurons were challenged with IL-1β (20 μg/ml), followed by polygodial (200 μM) and capsaicin (10 μM). Cells were washed with HBSS before application of each new agonist. Individual neurons were identified and analyzed for fluorescence intensity changes offline.

### Whole-cell patch-clamp recordings

Whole-cell patch-clamp recordings were performed as described previously^27^. Nodose ganglia neurons from Vglut2-GCaMP3 mice were selected for recording by their calcium fluorescence response to batch application of IL-1β (20 μg/ml). For patch-clamp recordings, neurons were visualized on a SliceScope system (Scientifica) with an Olympus BX51 microscope and Luigs and Neumann micromanipulators. Glass electrodes had a resistance of 2-4 MΩ and contained an intracellular solution (in mM): KCl 140, HEPES 10, EGTA 5, Mg-ATP 2, NaGTP 0.3, MgCl_2_ 2, phosphocreatine 10, pH 7.25 adjusted with KOH. The cells were perfused at a rate of 2 ml/min at room temperature (20-22°C) with an external bath solution containing the following (in mM): 140 NaCl, 5 KCl, 10 HEPES, 10 D-glucose, 2 CaCl_2_, and 1 MgCl_2_, pH 7.3-7.4. Whole-cell currents were acquired using an Multiclamp 700B amplifier (Molecular Devices, Union City, CA) and Windows PC running pClamp 11 software (Molecular Devices). Recordings were filtered at 2 Hz and sampled at 10 Hz. Data was analyzed offline using Clampfit 11 (Molecular Devices) and Origin 2019.

### Recording procedure and analysis

Mice were induced with general anesthesia using isoflurane at 2.5% in 100% oxygen at a flow rate of 1 L/min, and maintained in supine position at 1.25% isoflurane during recording. The left vagus nerve was surgically exposed and placed on electrode as described previously^26^. The electrophysiological signals were digitized from the vagus nerve using a Plexon data acquisition system (Omniplex, Plexon Inc., Dallas, Texas) as described previously^26^. Recordings were sampled at 40 kHz with a 120 Hz filter and 50 gain. All signals were recorded from either wild-type or TRPA1-deficient mice using a bipolar cuff electrode referenced to the animal ground placed between the right salivary gland and the skin. Following acquisition of the baseline activity (10 min), 350ng/kg recombinant human IL-1β (eBioscience, San Diego, California) was administered intraperitoneally; recordings were then continued for 10 min post-injection. Analysis of the neural recordings was done on Spike2 software (version 7, CED) as described previously^26^.

### Cytokine analysis

Mice were challenged intraperitoneally with 3.2 μg/kg IL-1β or saline (200 μl). Blood and spleen were collected 3 hr later for cytokine analysis. The spleen was homogenized using a bullet blender homogenizer (Next Advance, Averill Park, NY, USA), the supernatants collected and stored at - 20 °C. Splenic protein level was measured using the Bradford assay. Levels of IL-6 and CXCL1 for both serum and spleen were measured using a custom mouse inflammatory electro-chemiluminescent kit (Meso Scale Discovery, Gaithersburg, MD, USA) according to manufacturer’s recommendations.

### Telemetry system for temperature recordings

Mice were induced with general anesthesia using isoflurane at 2.5% in 100% oxygen at a flow rate of 1 L/min and maintained in supine position at 2.0% isoflurane. A midline incision was made, and an ETA-F10 temperature implant (DSI New Brighton, MN) was placed in the peritoneal cavity tacked to the peritoneal wall. After 5 days of recovery period, the mice were placed onto the DSI receiver. Baseline body temperature was recorded for 1 hour prior to IL-1β or saline injection. The recording was continued for 5 hr post-IL-1β administration.

### Cecil ligation and puncture

A standardized model of cecal ligation and puncture (CLP)–induced sublethal polymicrobial sepsis was used, as previously described^28^. Mice were anesthetized using ketamine 100 mg/kg and xylazine 8 mg/kg, administered intramuscularly. The cecum was isolated and ligated below the ileocecal valve and then punctured with a 22-gauge needle. Following approximately 2 mm of stool extrusion, the cecum was returned to the abdominal cavity. The abdomen was closed with surgical clips. An antibiotic, Primaxin (Imipenem-Cilastatin, 0.5 mg/kg, subcutaneously, in a total volume of 0.5 ml/mouse) was administered immediately after CLP as part of the resuscitation fluid. Mice were monitored for survival and sepsis-associated clinical signs. Disease severity scoring was done on days 3 and 6 post CLP using the murine sepsis score^29^.

### Statistical analysis

All statistical tests were performed with Graph Pad Prism 6 software (Graphpad, La Jolla, CA). Values are presented as individual samples and or mean ± SEM. Statistical analysis of mean differences between groups was performed using two-way ANOVA, paired t-test, Mann-Whitney U and Long-rank tests as indicated in respective results. All P values and n values are indicated in figure legends. P values equal to or less than 0.05 were considered significant.

## Data availability

The data that support the findings of this study are available from the corresponding authors upon reasonable request.

## ACKNOWLEDGMENTS

This study was supported by grants from the National Institute of Health (NIGMS 1R35GM118182-01 to K.J.T., and NIAID 1P01AI102852-01A1 to K.J.T. and S.S.C.).

## AUTHOR CONTRIBUTIONS

H.A.S., E.H.C., S.S.C. and K.J.T. designed research; H.A.S., M.G., Q.C., A.S., M.E.A., A.M.K., J.H.L., T.T., and T.S.H. performed research; H.A.S., M.G., E.H.C., Q.C., A.S., S.S.C. and K.J.T. analyzed and interpreted data; H.A.S., M.G., E.H.C., U.A., S.S.C. and K.J.T. wrote the manuscript; V.A.P. provided additional comments and contributed to finalizing the manuscript.

## MATERIALS and CORRESPONDENCE

Correspondence and requests for materials should be addressed to S.S.C. or K.J.T.

---

**Supplemental Data Fig. 1.**
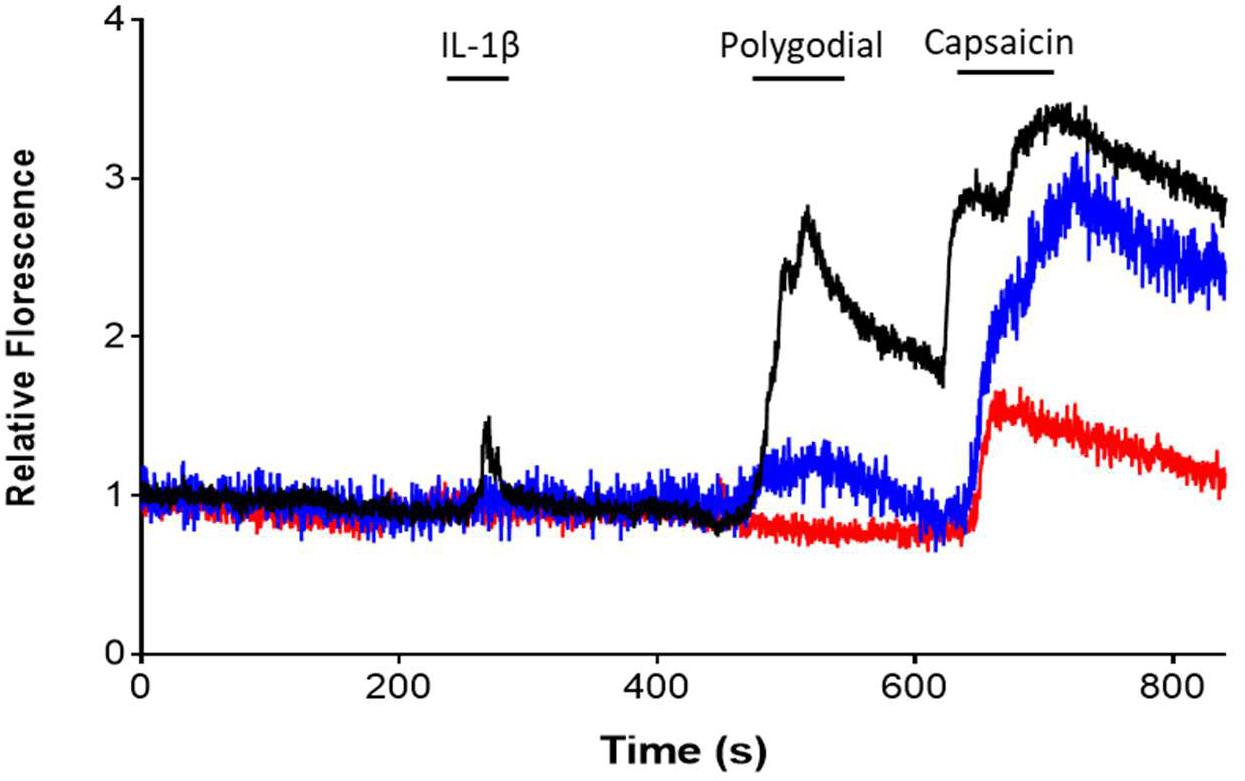
IL-1β induces activation of nodose sensory neurons responsive to polygodial and capsaicin. Representative GCaMP3 fluorescence signal in whole mount nodose ganglia showing responses to IL-1β. Polygodial (10 μM) and capsaicin (10 μM) are applied to identify TRPA1- and TRPV1-expressing neurons, respectively. Examples of neurons responding to IL-1β, polygodial and capsaicin (black line), to polygodial and capsaicin (blue line), and only capsaicin (red line) are shown.

**Supplemental Data Fig. 2.**
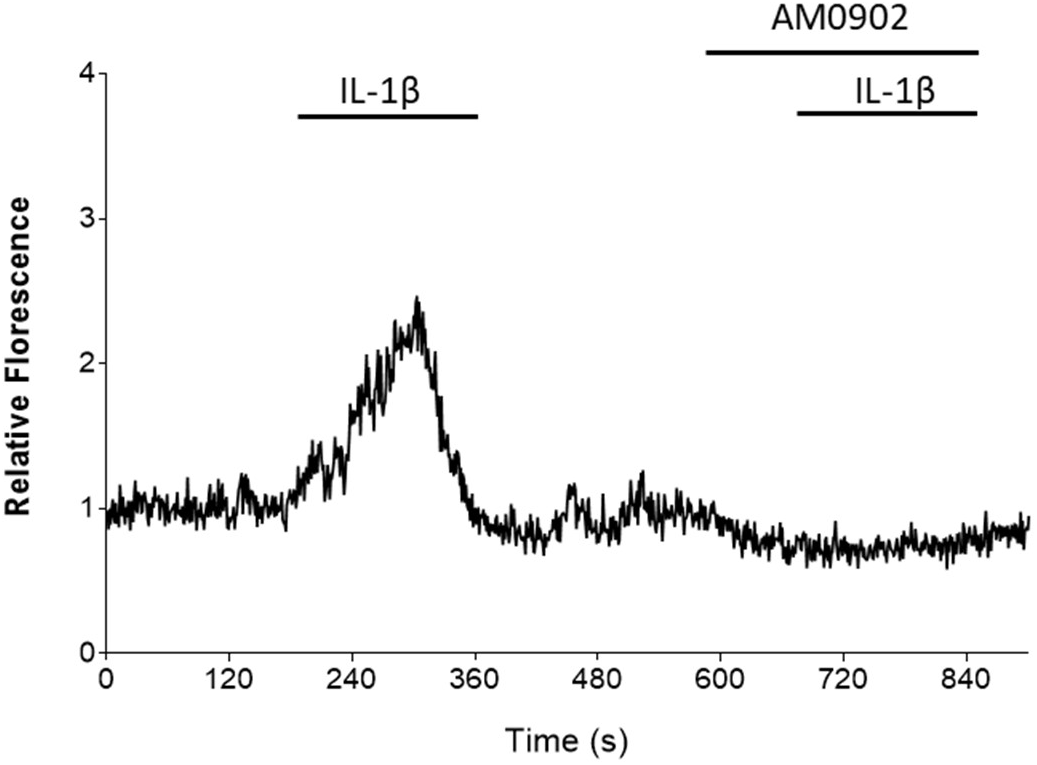
TRPA1 is required for IL-1β-mediated activation of nodose sensory neurons. Representative GCaMP3 fluorescence signal in cultured nodose ganglia neurons from Vglut-GCamp3 mice showing responses to IL-1β (20 μg/ml) in the absence and presence of TRPA1 antagonist AM0902 (10 μM).

**Supplemental Data Fig. 3.**
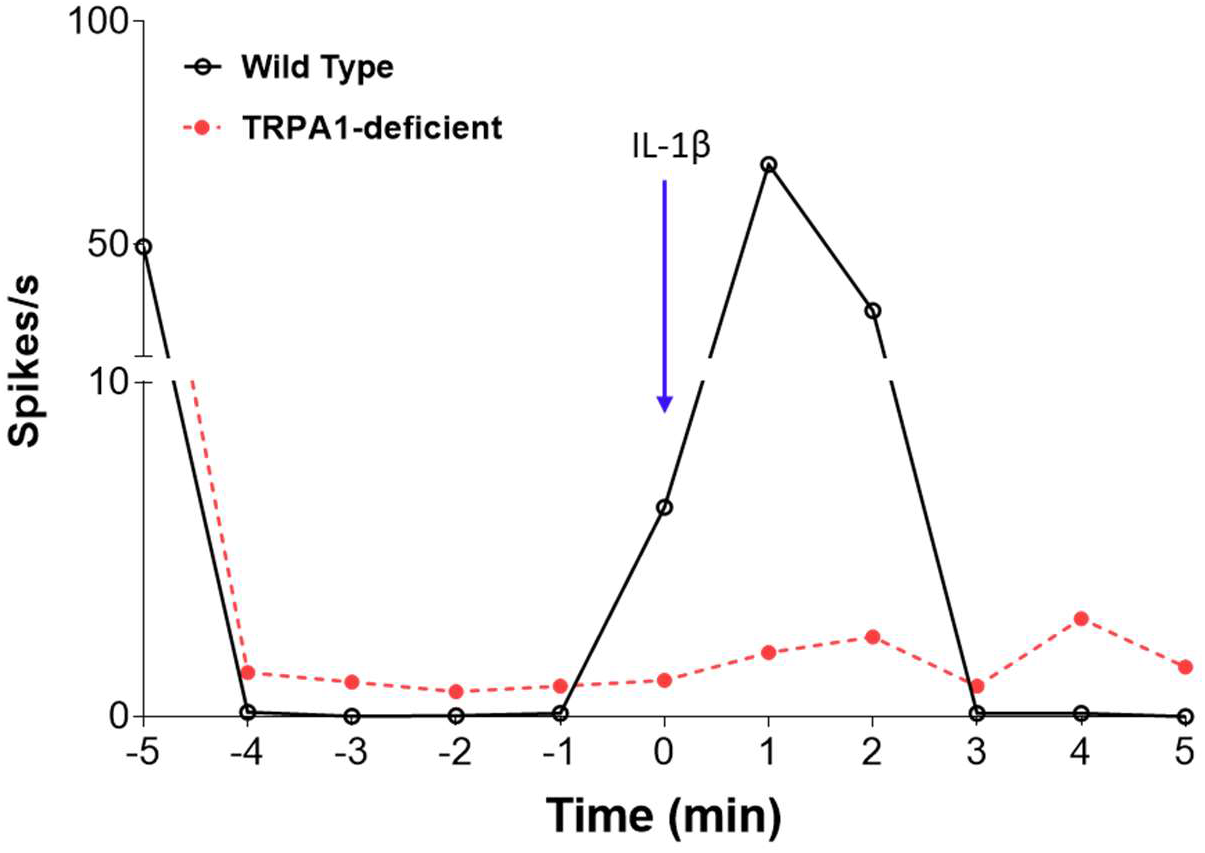
IL-1β fails to induce vagus nerve activation in TRPA1-deficient mice. Example of the effect of IL-1β administration (350 ng/kg, administered at time 0 min) on vagus nerve activity in wild-type (black line) and TRPA1-deficient mice (red dashed line). Representative spike rate in 1 min bins for the 5 min pre and post IL-1β administration is show

**Supplemental Data Fig. 4.**
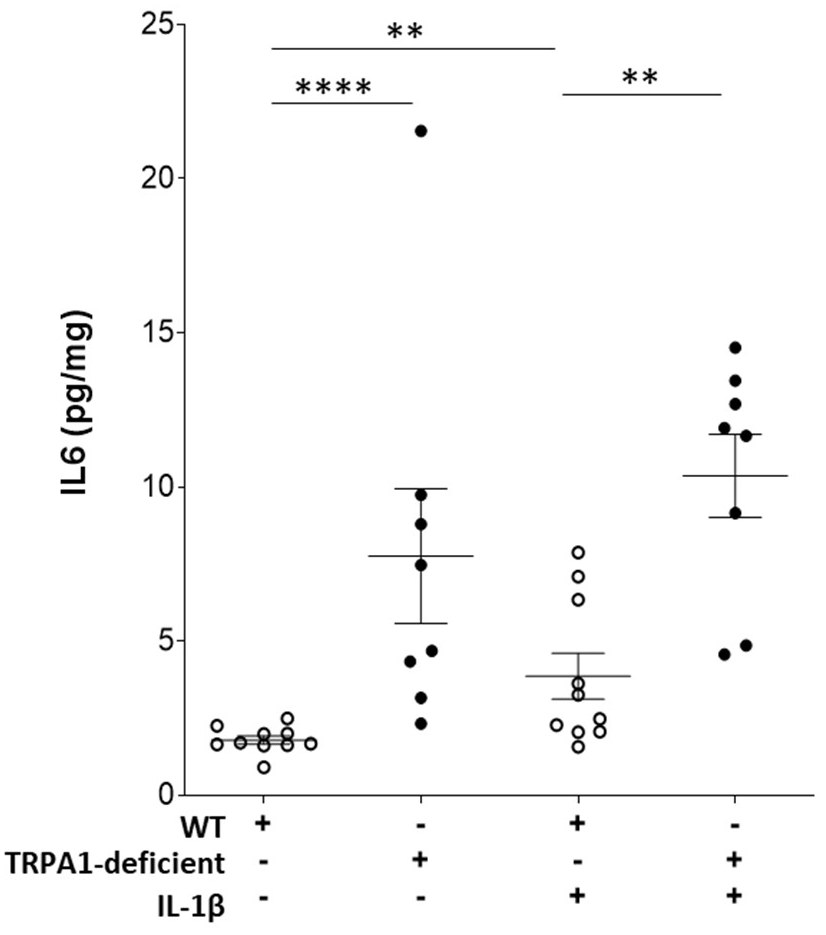
Increased splenic IL-6 levels in TRPA1-deficient mice. Splenic levels of IL-6 in wild-type and TRPA1-deficient mice injected intraperitoneally with either saline (n = 8-10 per group) or IL-1β (3.2 μg/kg, n = 8-10 per group). Mean ± SEM. **p<0.01, ****p<0.0001 (Mann-Whitney test).

**Supplemental Data Fig. 5.**
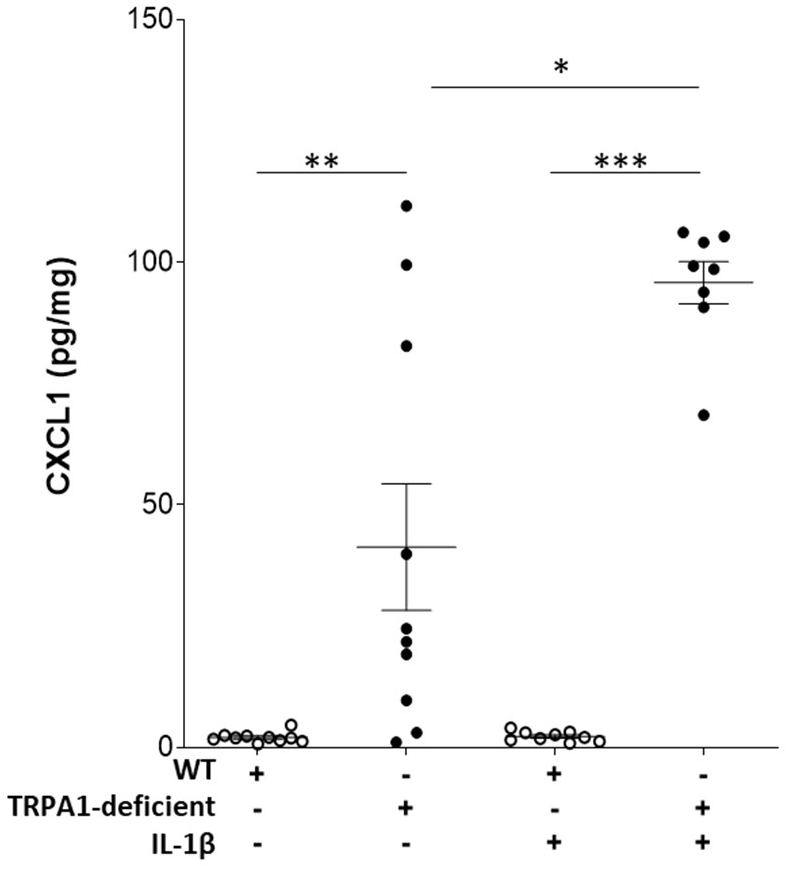
Increased splenic CXCL-1 levels in TRPA1-deficient mice. Splenic levels of CXCL-1 in wild-type and TRPA1-deficient mice injected intraperitoneally with either saline (n = 10 per group) or IL-1β (3.2 μg/kg, n = 8-9 per group). Mean ± SEM. *p<0.05, **p<0.01, ****p<0.0001 (Mann-Whitney test).

**Supplemental Data Fig. 6.**
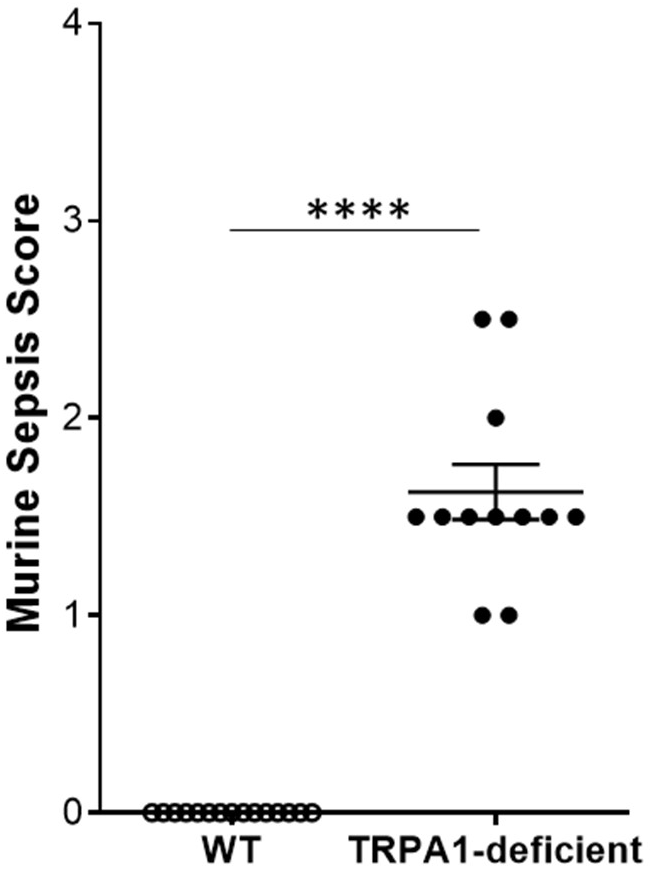
Increased sepsis severity in TRPA1-deficient mice. Murine Sepsis Score on post-CLP sepsis day 6 in wild-type (n = 15) and TRPA1-deficient mice (n = 12). Mean ± SEM. ****p<0.0001 (Mann-Whitney test) comparison between wild-type and TRPA

